# Invariant activity sequences across the mouse brain

**DOI:** 10.64898/2025.12.20.695676

**Authors:** Célian Bimbard, Kenneth D Harris, Matteo Carandini

## Abstract

Given the variety and plasticity of their circuits, different brain regions could potentially produce profoundly diverse patterns of population activity. Alternatively, the population activity could exhibit canonical patterns that are similar across the brain and stable over time. Here we show that a pattern observed in the cerebral cortex, where neurons fire in 10-100 ms sequences, is stable across weeks and ubiquitous across the brain. In mouse visual cortex, neurons respond to visual stimuli with fixed latency, forming a sequence. Similar sequences appeared during spontaneous activity: each neuron was coupled to the population with a characteristic strength and delay, which remained stable for weeks. Brainwide recordings revealed such sequences in every brain region. These results reveal a stable scaffold for neural activity across the brain.

## Introduction

The different regions of the brain contain radically different circuits and cell types, and are subject to multiple forms of plasticity. As a result, the populations of neurons in different brain regions could in principle exhibit profoundly diverse and mutable patterns of activity. Alternatively, the population activity could exhibit patterns that are canonical: ubiquitous across regions and stable over time.

Activity patterns that might potentially be canonical occur in the cerebral cortex, where neurons fire in 10-100 ms sequences. Some cortical neurons (“leaders”) tend to fire before their peers (*1*) while others (“followers”) fire after (*2*), forming sequences lasting 10-100 ms. Such sequences occur not only in the rodent cortex – where they are seen in somatosensory (*3, 4*), auditory (*4, 5*), visual (*6-8*), and prefrontal (*9*) areas – but also in the 3-layered cortex of turtles (*2*), the visual cortex of marmosets (*7*), and the temporal cortex of humans (*10-12*). The rodent auditory cortex, moreover, exhibits similar sequences spontaneously and following sound onsets (*4*). Likewise, the human temporal cortex exhibits the same sequences during a task and spontaneously (*10, 11*).

We hypothesized that these sequences of activity may form a stable scaffold for population activity in all brain regions. This hypothesis makes three predictions. First, the sequences should be invariant over the large range of firing rates that neurons produce, depending on the strength of their drive. Second, the sequences should be invariant over time, with the same neurons firing in the same order over multiple days. Third, and crucially, the sequences should be present in all brain regions, not only in the cerebral cortex.

Here we verified all three of these predictions. Chronic Neuropixels recordings in the visual cortex revealed that the sequences interact multiplicatively with the firing rate code used to represent stimuli and even occur spontaneously, thus being largely invariant to the range of firing rates produced by the neurons. The same recordings revealed that the sequences remain fixed over weeks, showing more stability across time than the firing rate code. Finally, acute Neuropixels recordings across the entire mouse brain revealed that the sequences of activity are ubiquitous, appearing in every brain region. These results reveal that sequences form a stable scaffold for neural activity across the brain.

## Results

We first focused on the visual cortex, where we could readily drive neuronal populations through visual stimulation. We inserted Neuropixels 2.0 probes (*14, 15*) chronically (*16*) in the visual cortex (VIS) of mice, and recorded from hundreds of neurons over several weeks (**Fig. 1**A). On each recording day, we presented a battery of the same 112 contrast-normalized images.

**Fig. 1.**
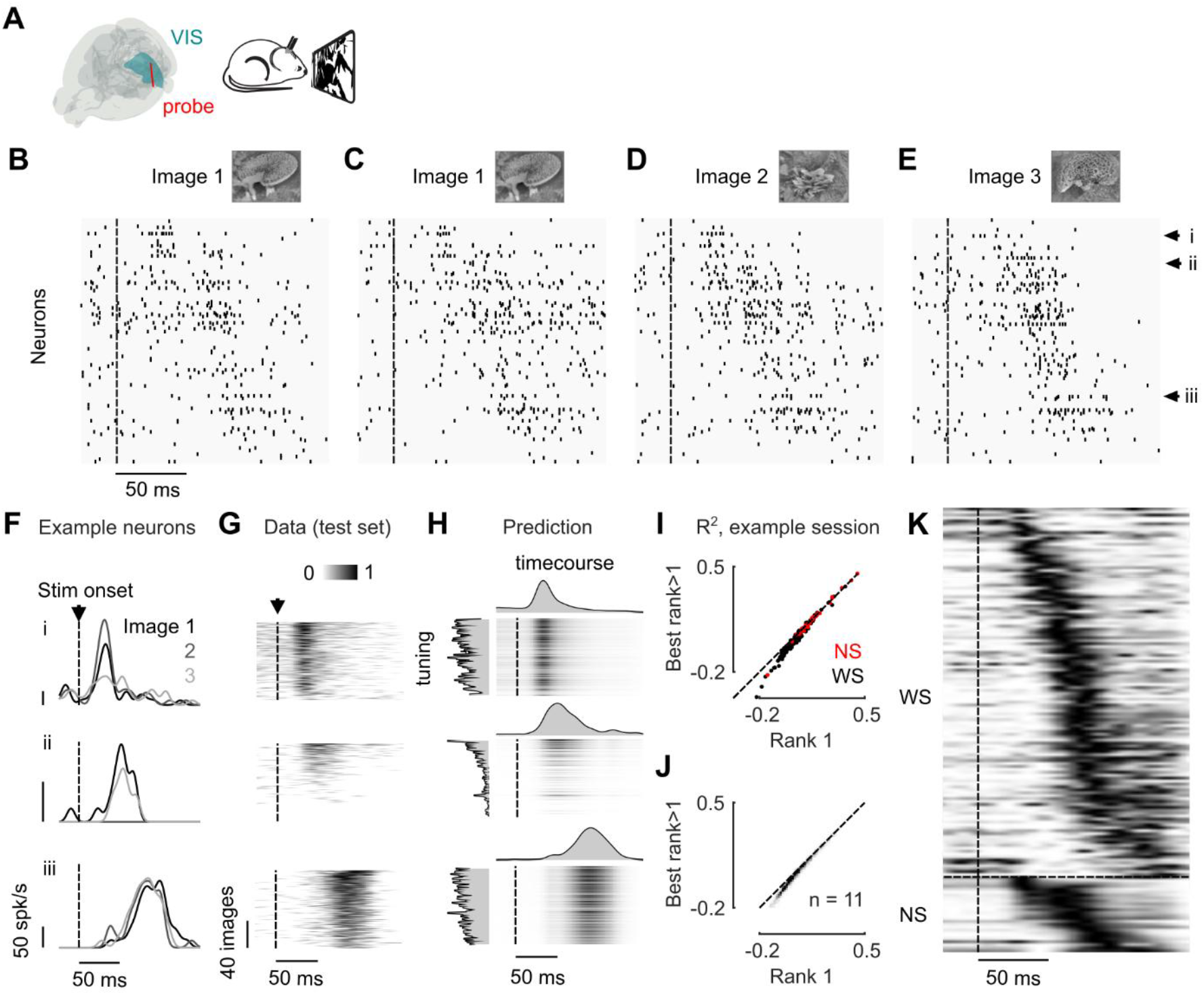
Stimulus-evoked activity sequences in visual cortex are invariant to firing rate, interacting multiplicatively with the responses. (**A**) Experimental setup for visual cortex (VIS) activity recordings and visual stimulation. (**B-E**) Raster for two example trials for image 1 (B, C) and one trial for two other images (D, E). Neurons were sorted by running Rastermap (*13*) on their average response to all images. (**F**) Peri-stimulus histograms (PSTHs) of 3 example neurons (i, ii and iii) in response to the same 3 example natural images as shown in (B)-(E) (shades of grey). The same neurons are indicated in (B-E) by the arrows. (**G**) PSTHs of the same 3 neurons in response to all images (normalized firing rate). Images were ordered by decreasing response amplitude for each neuron, measured on one half of the data (train set), while we display the responses on the other half (test set). (**H**) Prediction of the responses to all images on the test set for the same neurons as in (F) and (G). The prediction comes from a “rank-1” model, which multiplies a single time course (top of each heatmap) with a single tuning profile (left of each heatmap). (**I**) Quality of fit of a rank-1 model vs. the best model with a rank higher than 1 and up to 4 for all neurons within an example session. (**J**) Same as (I), for all neurons across all sessions of all animals.(**K**) Time courses obtained from the rank-1 model for all neurons, sorted by Rastermap (*13*), for the example session (normalized between 0 and 1 for each neuron).

The responses of individual VIS neurons to images had a characteristic time course, which formed a graded sequence largely conserved across stimuli. We used the sorting algorithm Rastermap (*13*) to order the neurons based on their time courses. The algorithm orders neurons by the similarity of their activity, revealing that the main organization of the responses is in the form of a sequence, with neurons firing early at the top and neurons firing late at the bottom (**Fig. 1**B to E). This sequence looked reliable across example single trials, and even across different images. For example, consider the activity of three example neurons exhibiting a range of latencies, in response to three example images (**Fig. 1**F). The amplitude of the responses differed across neurons: the first responded to images 1 and 2; the second to images 1 and 3, and the third to all three images. However, the time course of their responses seemed largely constant across images. The same results were obtained when looking at the responses to all images (**Fig. 1**G): the delay of the responses showed a strong variation across neurons but a small variation within neurons (comparison of variance of delays across neurons vs. across images in this example session: p < 0.001, n = 198 neurons and n = 112 images, two-sided Wilcoxon rank sum test.

The responses of each neuron across image stimuli could be well approximated as the product of a single time course and a single tuning profile. For each neuron, we applied non-negative matrix factorization to the matrix of responses as a function of time and image using one factor to obtain the best time course (one value at each time point) and the best tuning profile (one value for each image). The product of the two (a rank-1 approximation) provided a good summary of the original responses (**Fig. 1**G and H). Indeed, cross-validation revealed that the rank-1 approximation was superior to models that used more factors in 87 ± 2 % of the recorded neurons across all 11 mice (mean ± s.e., **Fig. 1**I and J). The time courses of each neuron are thus largely fixed across stimuli, and they gate *(4, 5)* the firing rate responses multiplicatively.

Across neurons, these time courses formed a sequence spanning 10-100 ms after stimulus onset (**Fig. 1**K). This large variation of time courses across neurons was true both for Wide Spiking neurons (WS), which are largely excitatory, and for Narrow Spiking neurons (NS), which are presumed inhibitory (**Fig. 1**K), as both WS and NS spanned the whole sequence.

This sequence proved to be a remarkably stable property of the neuronal population, remaining constant over weeks. We tracked the same cells across our chronic recordings for weeks using UnitMatch (*17*) (**Fig. 2**A), and observed that the sequence obtained on one day was essentially unchanged the next day (**Fig. 2**B and C). This level of stability was observed over weeks, with minimal signs of change over 30 days of recordings (slope = -0.002 ± 0.001, mean ± s.e., p = 0.16, n=4 mice, t-test, **Fig. 2**D). In all 4 mice, the neurons’ time courses were more stable *(8)* than their tuning, which showed a larger change over those 30 days (slope = -0.006 ± 0.002, mean ± s.e., p = 0.07, n = 4 mice t-test; comparison, p = 0.11, n = 4, paired t-test, **Fig. 2**E). This change in tuning is consistent with reports of drifts in the visual representation (*18-20*) (but see Refs. (*21-24*)).

**Fig. 2.**
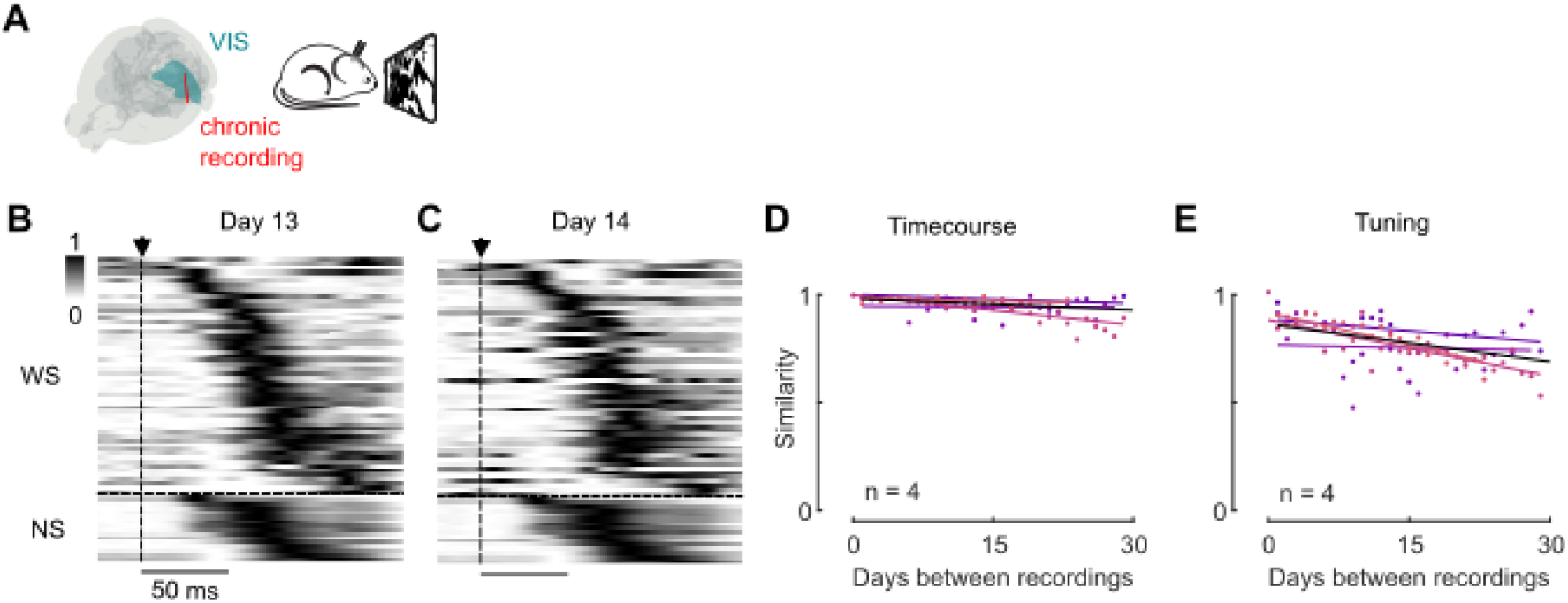
Stimulus-evoked activity sequences in visual cortex are invariant over time, remaining stable across weeks. (**A**) Experimental setup for chronic VIS activity recordings. (**B**) Time courses obtained from the rank-1 model for all neurons, sorted by Rastermap (*13*), for the example session on day 13 (normalized between 0 and 1 for each neuron). (**C**) Same as (M), but during the following day (day 14), after tracking the same neurons across both sessions. (**D**) Similarity (noise-corrected correlation) of the neurons’ time courses across recordings for different mice (*colours*) and the average (*black*). (**E**) Similarity of the neurons’ tuning profiles across recordings.

We then asked whether we could find similar sequential structure in spontaneous activity. To do so, we considered the coupling of each individual neuron to the summed activity of all neurons (the population rate; **Fig. 3**B). Specifically, we computed the spike-triggered population rate (*25*) for each neuron, which we refer to as “population coupling” (**Fig. 3**C). The strength of this coupling at zero delay is a primary determinant of the pairwise correlations in a population (*25*) and (unlike correlation (*26, 27*)) is not affected by the neuron’s firing rate (*25*). As expected (*25, 28*), the strength of population coupling exhibited a large variation across neurons, with some neurons (“choristers”) highly coupled to the population and others (“soloists”) barely coupled (**Fig. 3**Cand E). In addition, we measured the delay of each neuron to population activity by computing the centre of mass of the population coupling profile; this is negative for “leader” (*1*) neurons which fire earlier than the majority of their neighbours, and positive for “follower” (*2*) neurons that fire after their neighbours. Neurons varied in both measures (**Fig. 3**C and H), with no hint of relationship between the two: the correlation between coupling strength and delay was 0.04 ± 0.05 in WS cells, and 0.08 ± 0.15 in NS cells (mean ± s.e., p = 0.83 for WS cells, p = 0.64 for NS cells, n = 11 mice, two-sided Wilcoxon rank sum test, **Fig. 3**D). The two measures thus define largely independent axes to describe population structure.

**Fig. 3.**
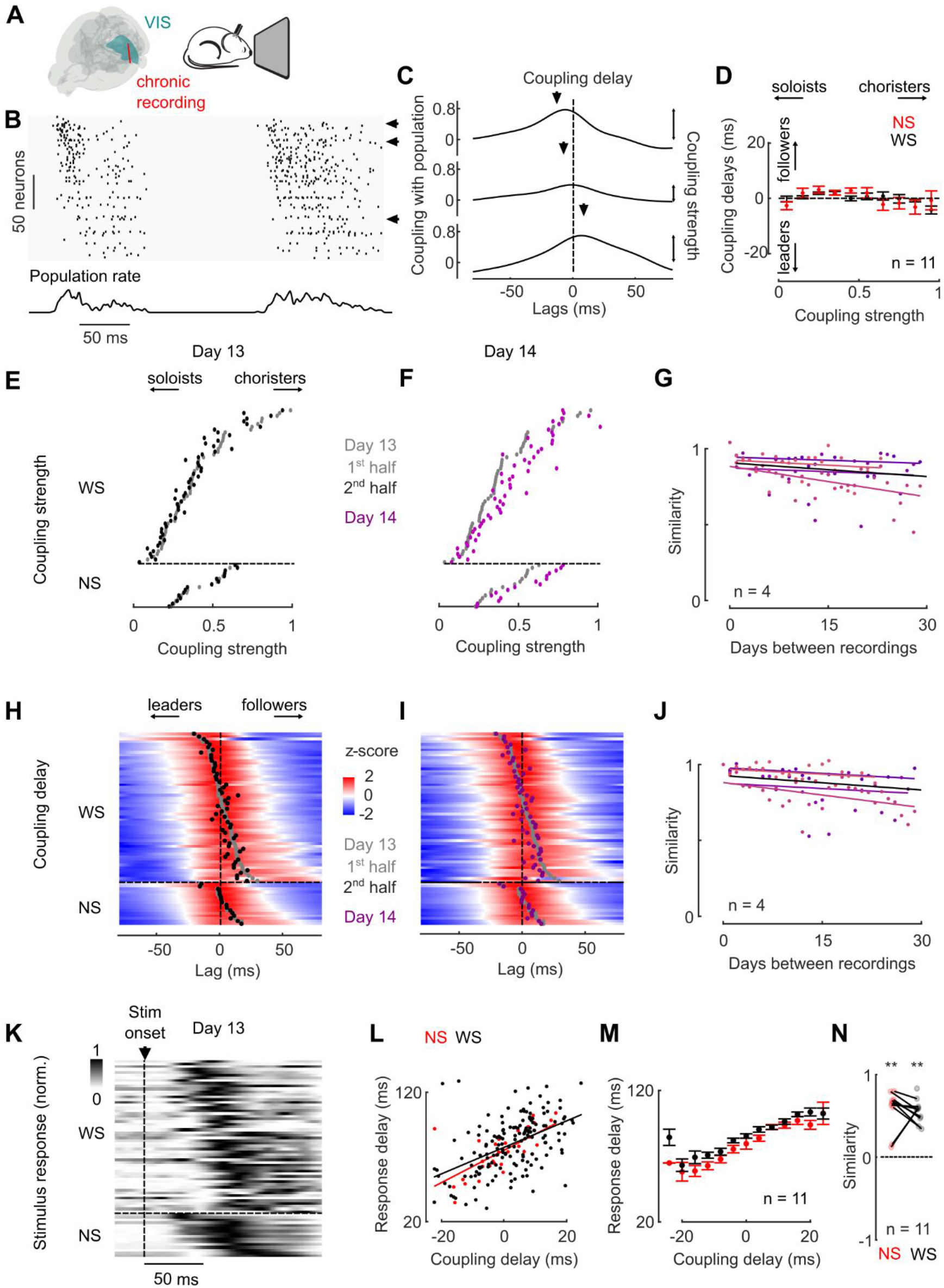
Spontaneous activity sequences in visual cortex are stable across days and resemble evoked sequences. (**A**) Experimental setup for spontaneous activity recordings. (**B**) Example raster of spontaneous activity (top) and average population rate (bottom), showing two individual and consecutive bursts of sequential activity. Neurons were ordered based on their population coupling delay. (**C**) Coupling with the population for 3 example neurons, designed by the black arrows in (B). Two properties were extracted: the coupling strength, i.e. the value of the coupling at time 0, and the coupling delay, i.e. the centre of mass of the population coupling. Note that our convention for the lag of the population coupling means that a negative delay corresponds to a neuron firing before the population. (**D**) Relationship between the coupling strength and the coupling delay of WS (*black*) and NS (*red*) neurons, averaged across all mice (n = 11, mean ± s.e.). (**E**) Coupling strength across all neurons for an example session recorded on example day 13. Neurons were ordered by decreasing coupling strength and separated according to their spike waveform (WS: wide spiking; NS: narrow spiking). Within days, the coupling strength for each neuron was computed for both the 1^st^ (*grey dots*) and the 2^nd^ (*black dots*) halves, showing high reliability. (**F**) Same as (E), on the next day, after tracking the same neurons across recordings. A high reliability was also observed between the two days (*grey dots:* day 1, 1^st^ half; *purple dots*: day 2, average over both halves). (**G**) Similarity of the neurons’ coupling strength across recordings. Different colours represent different mice. (**H**)**-**(**J**). Same as (E)-(G), but for coupling delays. Population couplings were z-scored for each neuron and ordered by their coupling delay during the 1^st^ half of the session. (**K**) Stimulus-evoked time courses of the same neurons, ordered as in (**H**) and (**I**) by delay during spontaneous sequences (normalized between 0 and 1 for each neuron). (**L**) Comparison of the spontaneous coupling delays and evoked responses delays, for both the putative excitatory (*black*, WS) and inhibitory (*red*, NS) neurons, in an example session. (**M**) Same as (L), with the average across all mice (n = 11, mean ± s.e.). (**N**) Similarity of the evoked and spontaneous delays across all animals (n = 11).

Thanks to the variation in coupling delay, the spontaneous activity in the mouse visual cortex thus formed stereotyped sequences (**Fig. 3**H), similar to the sequences observed after eye movements and stimulus onsets (*6, 7*) and to the spontaneous sequences observed in other cortical regions in other species (*2-4, 10, 11*).

Both the magnitude and delay of population coupling were highly consistent over weeks. A cell’s coupling strength was largely constant across recording sessions (**Fig. 3**F), exhibiting a similarity that showed little sign of decay across weeks (slope = -0.003 ± 0.001, mean ± s.e., n = 4 mice, **Fig. 3**H): neurons that are choristers one day, remain so.

The sequences seen during spontaneous activity also remained stable over time. The spontaneous sequences measured on one day were essentially identical the next day (**Fig. 3**H and I), with coupling delay showing a similarity that showed little sign of decay over weeks (slope = -0.003 ± 0.001, mean ± s.e., n = 4 mice, **Fig. 3**J).

Moreover, the sequences seen during spontaneous activity strongly resembled the sequences evoked by visual stimuli. To illustrate the similarity, we reordered the sequence we had seen in response to images (**Fig. 2**B) so that neurons have the same ordering as the spontaneous sequences (**Fig. 3**K). The spontaneous and driven sequences are clearly similar, and indeed the response delay correlated strongly with the coupling delay measured during spontaneous activity in this example session (r = 0.53) (**Fig. 3**L) and across all sessions and animals (**Fig. 3**M). We computed the similarity of the two (Methods) and found it to be significant for both WS and NS neurons (WS: similarity s = 0.58 ± 0.18, p = 0.002; NS: s = 0.66 ± 0.18, p < 0.001, n = 11 mice; median ± mad, two-sided Wilcoxon signed-rank test), with no significant difference between the two (p = 0.43, paired two-sided Wilcoxon signed-rank test) (**Fig. 3**N).

We next asked whether similar sequential patterns of activity occur also in other brain regions. We did so using acute recordings covering an entire hemisphere of the mouse brain, which we have performed with the International Brain Laboratory (*29*). We examined the activity of 54,553 neurons in 230 brain regions, obtained from 419 recordings involving 644 insertions of Neuropixels 1.0 and 2.0 probes (*14, 15*). We further selected recordings with more than 5 neurons. We focused on periods of spontaneous activity (recorded after the mice had performed the IBL task) and performed the same analysis that we had performed on the spontaneous activity of the primary visual cortex (**Fig. 4**A). Because the electrophysiological signatures of different cell types are generally less understood outside the cortex, in these brainwide recordings we did not attempt to separate putative excitatory and inhibitory neurons.

**Fig. 4.**
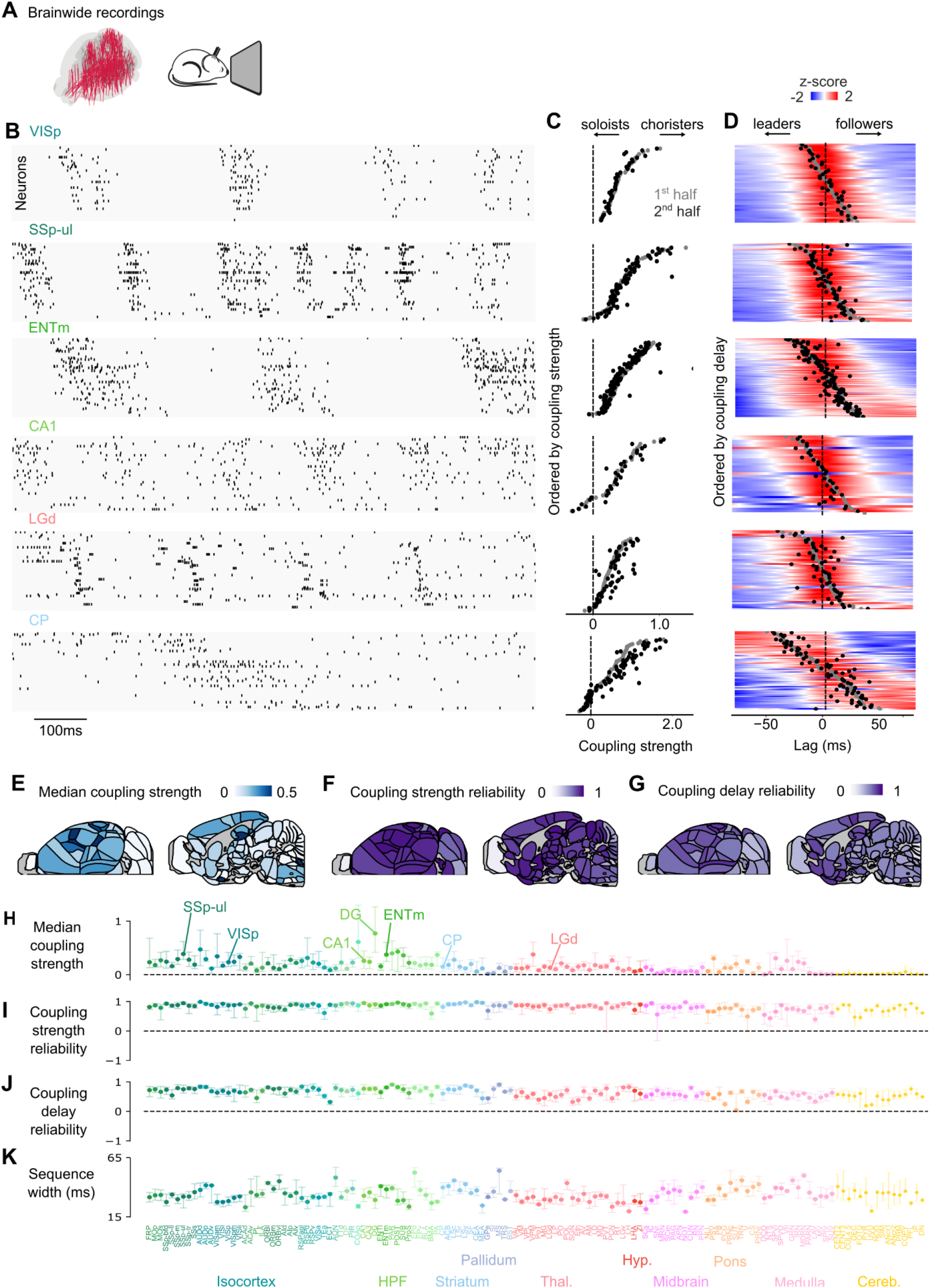
Spontaneous activity sequences are found across the entire mouse brain. (**A**). Brainwide recordings from the International Brain Laboratory brainwide map dataset (*29*). (**B**) Raster plots of 1s-snippets of activity for 6 example brain areas, cortical and subcortical: primary visual cortex (VISp), the upper limb field in primary somatosensory area (SSp-ul), medial entorhinal cortex (ENTm), hippocampal field CA1, dorsal lateral geniculate nucleus (LGd) and caudate-putamen (CP). Neurons were ordered according to their coupling delay on the first half of the session. (**C**) Coupling strength for the same areas, shown for both the first (*grey*) and the second (*black*) half of each session. Neurons were reordered according to their coupling strength on the first half of the session. (**D**) Z-scored population coupling for the same brain regions as in (B), showing the coupling delays, and ordered by coupling delays. (**E**) Median coupling strength across neurons for every recorded area of the brain. (**F**) Same as (**E**), for the coupling strength reliability. (**G**) Same as (**E**), for the coupling delay reliability. (**H**) Median coupling strength across neurons, across all 138 recorded brain regions with at least 5 recordings. The central dot shows the median across recordings, and the error bars the 25^th^ and 75^th^ percentiles. (**I**) Same as (H), with the coupling strength reliability within days. (**J**) Same as (H), with the coupling delay reliability within days. (**K**) Same as (H), with the sequences’ width.

Sequences of activation appeared in essentially every brain region. For every region, we measured the strength and delay of the coupling of each neuron’s activity to the population of neurons in the same region, assessing their reliability by correlating the measures obtained in two separate halves of a given recording. When inspecting ongoing activity and ordering neurons by their coupling delays, clear sequences were visible, with the activity of “leader” neurons preceding that of “follower” neurons (**Fig. 4**B). Over the whole session, neurons in essentially all brain regions showed reliable and diverse strengths of population coupling, with “soloist” neurons showing low coupling and “chorister” neurons showing strong coupling (**Fig. 4**C). Furthermore, the individual sequences observed in the rasters of ongoing activity (**Fig. 4**B) were well captured by their delayed population couplings (**Fig. 4**D). As examples, we illustrate these results both in areas of the cerebral cortex where the sequences have already been observed, such as the primary visual (*7*) and somatosensory (*3, 4*) (VISp and SSp) areas and in additional regions such as entorhinal cortex (ENTm) and subcortically in hippocampus (CA1), visual thalamus (LGd), and the striatum (CP).

To verify the robustness of these findings, we computed the population coupling strength and delay for all the recorded brain regions. Coupling magnitude was uneven across brain regions (p < 0.001, Kruskal Wallis test). In some regions it was stronger than in the primary sensory areas, e.g. in the dentate gyrus (DG) and in the medial entorhinal cortex (ENTm), and in others it was weaker, e.g. in the cerebellum (**Fig. 4**E and H). Coupling magnitude and delay were however both highly reliable (consistent across two separate halves of a recording) in 138 out of the 138 brain regions (selected with at least 5 recordings, **Fig. 4**F, G, I and J-). The duration of the sequences also varied across regions (p < 0.001, Kruskal Wallis test). The range of coupling delays (measured between the 10^th^ and 90^th^ percentiles of the delay distribution) had an average of 35 ms and standard deviation of 6 ms across brain regions (n = 138, **Fig 4**K), but could be shorter in certain brain regions (e.g., VISp: 30 ± 8 ms; SSp-ul: 31 ± 10 ms, median ± m.a.d.) or longer in others (e.g., ENTm: 37 ± 8 ms; CP, 40 ± 10 ms).

The observed coupling strengths and delays in the cerebral cortex were not explained by straightforward, layer-based circuitry. It is known that neurons may fire at different times across cortical depths (*30, 31*). Here, coupling delays were largely heterogeneous even within layers (**Fig. S**1A to D). We quantified the amount of reliable variance in the distribution of strengths and delays and found that the layer position of the neurons explained <10% of the reliable variance, leaving 90±1% and 93±1% of the reliable variance unexplained for the coupling strengths and delays (**Fig. S**1E and F).

We finally asked whether the coupling (*28, 33*) and the sequences of activity that are observed across the brain can be coordinated with each other. Though simultaneous recordings were not the goal of the IBL brainwide map, each Neuropixels probe invariably recorded from more than one region, and each session typically involved recordings from two probes. We can thus ask if the coupling that we have observed within all brain regions can also occur across regions. To illustrate the approach, consider a session where primary visual cortex (VISp) was recorded at the same time as medial entorhinal cortex (ENTm, **Fig. 5**A and B). In this session, we can use the population activity in VISp as a “source” to measure population coupling not only in VISp itself (**Fig. 5**C) but also in ENTm (**Fig. 5**D), which are both “targets”. The same analysis can be run by using ENTm as the source, and measuring the coupling to this source population in both target regions (**Fig 5**E and F).

**Fig. 5.**
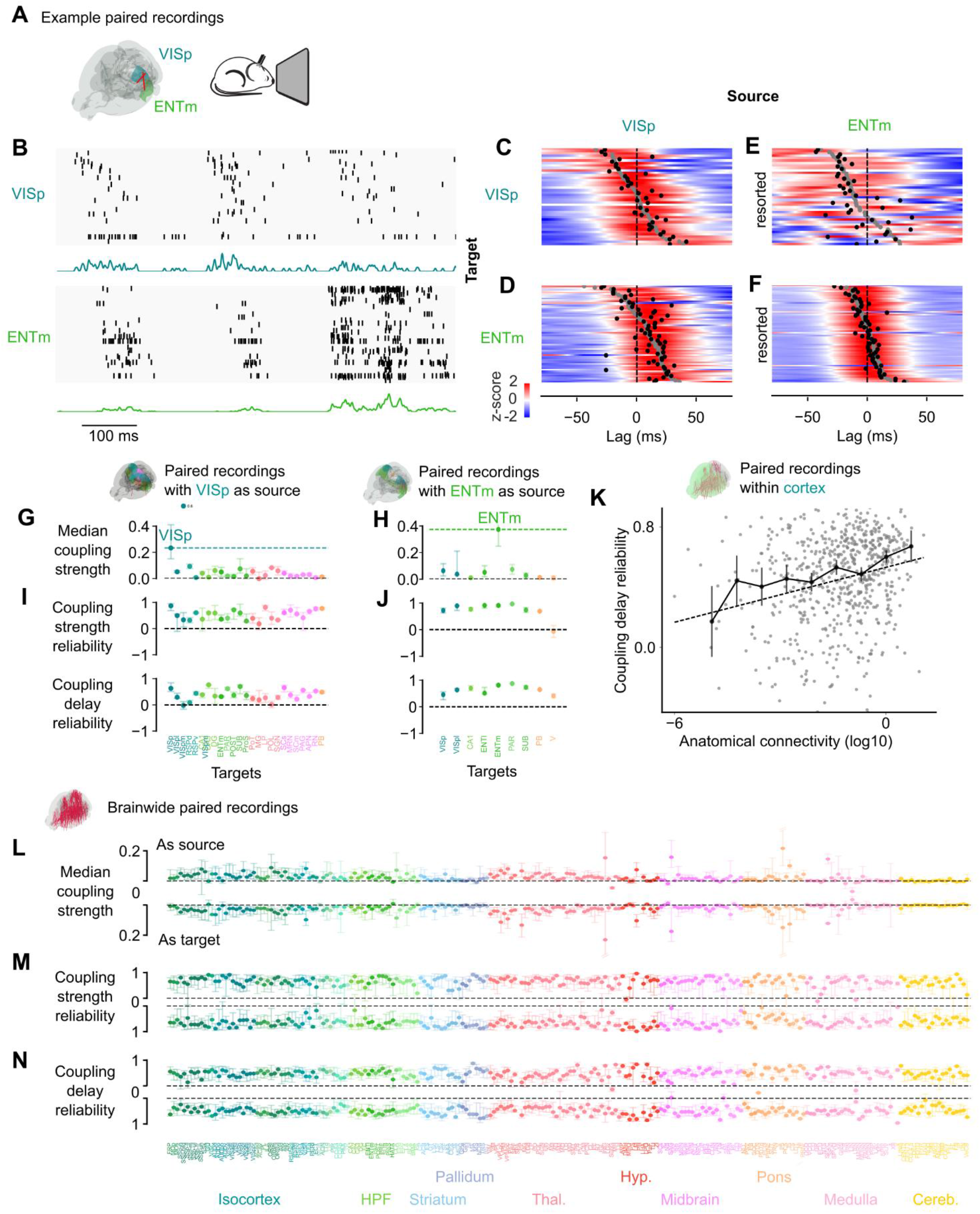
Activity sequences are occasionally coherent across brain areas. (**A**) Example paired recording. (**B**) Rasters of activity with individual neurons activity in VISp (top) and ENTm (bottom), and VISP and ENTm average population rates. Neurons were ordered according to their coupling delay with the population rate in VISp. (**C**) Z-scored coupling of VISp neurons with VISp population rate, with the coupling delays for the 1^st^ half (*grey dots*) and the 2^nd^ half (*black* dots) of the recordings. (**D**) Same as (C), with VISp as a source and ENTm as a target. (**E**) Same as (C), with ENTm as a source and VISp as a target. Neurons were resorted based on their coupling delay with ENTm. (**F**) Same as (E), with ENTm as a source and ENTm as a target. (**G**) Median coupling strength of each brain region paired with VISp. The dashed line shows the median coupling strength for VISp, the error bars indicate the 25^th^ and 75^th^ percentiles across recordings. (**H**) Same as (G), for each brain region paired with ENTm. (**I**) and (**J**) Same as (G) and (H) but showing the coupling strength reliability and the coupling delay reliability with VISp (I) and ENTm (J) as a source. (**K**) Relationship between the coupling delay reliability of each pair of recording regions with their anatomical connectivity, measured by the Allen Institute Connectivity Atlas (*32*). (**L-N**). Coupling delay reliability (L), coupling strength reliability (M) and median coupling strength (N) for each brain region as a source, averaged over all targets (top) or as a target, averaged over all sources (bottom). The dot represents the median across all recordings involving this area and error bars indicate the 25^th^ and 75^th^ percentiles.

The results of this analysis in VISp and ENTm revealed that neuronal coupling can extend beyond individual regions, and form reliable sequences of activation. Visual inspection of the simultaneous rasters occasionally revealed activation sequences coordinated across regions (**Fig 5**B). On average across the session, the activity in ENTm was indeed coupled to the activity in VISp, and formed a sequence that was synchronous to the one occurring there (**Fig. 5**D). Similarly, the activity in VISp was coupled to the activity in ENTm, and could form a sequence that was synchronous to the one occurring there (**Fig. 5**E). The sequences in each target region, however, showed some dependence on the source region (the sorting of neurons in **Fig. 5**C and D is somewhat different from the one in **Fig. 5**E and F), perhaps suggesting that the order of the sequence contains a signature of the region to which it is coupled.

These occasionally coordinated sequences with VISp and ENTm activity as sources could be observed across many target brain regions. For example, 24 regions (“targets”) were recorded simultaneously with VISp (“source”, **Fig. 5**G), and 8 regions were recorded simultaneously with ENTm (**Fig. 5**H). For each of these combinations of targets and source, we computed the coupling strengths and delays of the neurons in the target areas. The mean coupling strength was generally lower across regions than within a region (**Fig. 5**G and H), suggesting that sequences were primarily local to each brain region. Nonetheless, the coupling strengths and delays across regions were generally reliable: this was true in 24 out of 24 brain regions recorded with VISp, and 7 out of 8 with ENTm (**Fig. 5**I and J). Thus, sequences were occasionally coordinated across regions. In the cerebral cortex, this reliable coupling across regions correlated with the anatomical connectivity measured by the Allen Institute Connectivity Atlas (*32*) (Spearman correlation = 0.29 across 648 pairs of cortical areas, p < 0.0001, Mantel test, **Fig 5**K).

Overall, each brain region exhibited weak but reliable coupling with the regions that were recorded simultaneously with it, whether as a source or as a target. For each brain region (target), we computed the strength and reliability of its coupling with all other simultaneously recorded regions (sources). We then obtained two averages: one over sources and one over targets. The former measures the overall coupling of neurons in each target region with the population rate of source regions, and the latter measures the overall coupling of the population rate of each source region to neurons outside it. The strengths of the cross-area coupling showed some variation across regions (**Fig. 5**L), and was invariably lower than the strength of coupling within regions (ratio of median coupling strength across vs. within regions: 0.12 ± 0.10 as sources, 0.13 ± 0.11 as targets, median ± mad across all brain regions). This was true in 218 out of 230 brain regions as sources and as targets, confirming that sequences are mainly local. However, the cross-area coupling delay and coupling strength were remarkably reliable in practically all regions (at least 222 out of 230), regardless of whether they were taken as a source or as a target (**Fig. 5**M and N). These results show that practically all brain regions are effective sources and targets of weak but reliable, coordinated coupling.

## Discussion

We have shown that neural activity exhibits strong coupling and is organized in 10-100 ms sequences not only in some areas of the cerebral cortex but also in essentially every region of the mouse brain, displaying sequences that are coordinated mostly within regions and occasionally across regions. These findings in the mouse brain extend those previously been obtained in the mouse visual cortex (*6, 7*), and in the somatosensory (*3, 4*) and auditory (*4*) cortex of the rat. Given that sequences have also been observed in the cerebral cortex of reptiles (*2, 34*) and primates (*7*), including humans (*10, 11*), it is tempting to speculate that they might be ubiquitous, shaping the structure of brainwide activity in all vertebrates.

Claiming the existence of patterns in neural activity could be controversial if the claim is based on complex and potentially biased statistical measures (*35, 36*). To ensure robustness and transparency we adopted extremely simple statistical measures: for each neuron we computed the population coupling (*25*) (which is simply the spike-triggered population rate) and we measured its height at delay zero (yielding coupling strength) and its centre of mass (yielding coupling delay) (*3*), assessing reliability by cross-validation: dividing the data in two within a session (or where applicable across sessions), and correlating the results across the two halves.

The sequences that we have observed interacted multiplicatively with the firing rate code, constraining but not mandating or impeding the participation of individual neurons. Indeed, in the visual cortex the participation of individual neurons in an activity sequence elicited by an image depended on the tuning of those neurons to that image. However, if the tuning of the neurons is such that they respond to the image, then their response occurred at a specific time in the sequence, which was largely fixed for each neuron. Indeed, the variation in timing within neurons across images was much smaller than that across neurons. These results echo those seen in rodent auditory cortex (*4, 5*) but contrast those proposed for the human temporal cortex (*12*). More precisely, the sequences interacted multiplicatively with the firing rate code that describes the tuning of the neurons, so that at any given time and for any given image, the response of the neuron could be well approximated by the product of a tuning curve that depends on images but not time, and a response profile that depends on time but not on images.

By recording from the same neurons over time, we established that the sequences – both evoked and spontaneous – are stable over weeks. In our 4 chronically recorded animals, the sequences evoked by stimuli in visual cortex were stable (*8*). By contrast, consistent with some (*18-20, 37-41*) but not all (*21-24, 42, 43*) reports, the firing rate code representing visual stimuli appeared to show some drift. We moreover discovered that the strength and delay of each neuron’s coupling with the population was highly stable across weeks. This stability agrees with the view that, overall, the sequences constitute a hard-wired scaffold (*11*) for neural activity. However, it is also possible that dynamical processes may in some cases modify the sequences through learning or plasticity (*6*).

As had been seen in other cortical areas in rats (*3, 4*) and humans (*10, 11*), the sequences occurred not only during sensory-driven activity but also during spontaneous activity. The fact that both forms of activity are constrained to the same spatiotemporal patterns might help explain the similarity between spontaneous and driven activity that has been reported multiple times (e.g. Refs. (*4, 44-48*)). The occurrence of the sequences during spontaneous activity was also a welcome observation in that it allowed us to extend the analysis from a region where we could elicit activity at will – the visual cortex – to the rest of the brain, where activity is generally ongoing and less easily elicited.

The sequences we have observed across the brain were very similar to the 10-100 ms sequences measured in various cortical regions (*2-4, 6, 7, 10, 11, 34*), but their duration was on the short end: the median sequence duration averaged ~35 ms across regions. It is hard to know if this duration is significantly shorter than in previous studies, however, because those studies did not quantify the sequence duration.

These rapid sequences may bear some relation to the sequences observed in hippocampal “replay” (*49-52*) and “preplay” (*53-55*), which typically last hundreds of milliseconds. These longer sequences appear not only in hippocampus but also in visual (*52*) and prefrontal (*56*) cortex, so, just as with the rapid sequences studied here, they may also appear in other brain regions. Though the rapid sequences are much briefer than the “preplay” and “replay” patterns, they are on the same time scale as the theta cycle, and as such they may be related to the alternating “sweeps” observed in hippocampus and entorhinal cortex during the theta cycle and during REM sleep (*57*).

The rapid sequences we observed also appear to differ from other sequences observed in various neural systems (*58*), which are generally substantially longer. These include 0.5-1.0 s songs in the zebra finch’s premotor nucleus HVC (*59*); ~1 s sequences in ex vivo slices of mouse cortex (*59*); ~1.5 s “playback” patterns in prefrontal cortex (*60*); multi-second sequences in many brain regions (*61*) associated with arousal events (*62*); periodic multi-second sequences in medial entorhinal cortex (*63*); and multi-second sequences in the mouse visual cortex (*63*).

The circuit basis of the sequences is currently unknown, but may arise from simple and known factors. Sequences of various durations arise readily in silico (*34, 64*) and even in organoids (*65*). Suggested mechanisms generally involve a synaptic chain of activation (*34*) shaped by intrinsic excitability (*66*) and conduction delays (*67*). The occurrence of sequences may be shaped by brain states (*68, 69*).

Sequences of spiking activity have been suggested as potential building blocks of information (*1, 70*), controlling the precision of sensory processing (*7*) or creating sequences of motor actions (*66*). Moreover, the occasional coherence of the sequences across pairs of regions suggests that sequences of activity may be specifically triggered throughout larger brain networks. These broader sequential events could possibly reflect switchable communication routes across the brain. Furthermore, different sequences may be associated with different source regions, generating a specific temporal “fingerprint” within each target region. The functional role of brainwide sequences, however, remains unknown. Addressing these questions will likely require new datasets obtained with massively parallel brainwide recordings.

## Acknowledgements

We thank Anna Lebedeva, Michael Okun, Enny van Beest, and Agnès Landemard for help with data and analysis, and Amirreza Bahramani for comments on the manuscript. This research was funded by the Wellcome Trust (Investigator Award 223144/Z/21/Z to MC and KDH, and Early Career Award 227065 to CB) and the Simons Foundation Collaboration on the Global Brain (award AN-NC-GB-IBL-00002672-16 to MC). MC holds the GlaxoSmithKline / Fight for Sight Chair in Visual Neuroscience.

## Author contributions

Conceptualization, funding acquisition, resources, and writing—review and editing: C.B., K.D.H. and M.C.; Methodology, software, formal analysis, and investigation: C.B.; Visualization and writing— original draft: C.B. and M.C.; Supervision: M.C.

## Data availability

Part of the data is already available online (*15*) (https://doi.org/10.5522/04/24411841), and the rest will be made available upon publication. Instructions for downloading the IBL data are available online (https://int-brain-lab.github.io/iblenv/notebooks_external/data_release_brainwidemap.html). The data can also be browsed online at the IBL website (https://viz.internationalbrainlab.org).

## Methods

Our study is based on two main datasets: one involving chronic recordings in the mouse visual cortex, performed in our laboratory, and one involving acute brainwide recordings, performed in collaboration with the International Brain Laboratory (*29*).

### Chronic recordings in visual cortex

Experimental procedures at UCL were conducted according to the UK Animals Scientific Procedures Act (1986) and under personal and project licenses released by the Home Office following appropriate ethics review.

### Surgeries

A brief (~1 h) initial surgery was performed under isoflurane (1–3% in O_2_) anaesthesia to implant a headplate made of titanium (~25 × 3 × 0.5 mm, 0.2 g) or steel (~25 × 5 × 1 mm, 0.5 g). In brief, the dorsal surface of the skull was cleared of skin and periosteum. A thin layer of cyanoacrylate was applied to the skull and allowed to dry. Thin layers of ultraviolet (UV)-curing optical glue (Norland Optical Adhesives #81, Norland Products) were applied and cured until the exposed skull was covered. The head plate was attached to the skull over the interparietal bone with Super-Bond polymer. After recovery, mice were treated with carprofen or meloxicam for 3 days, then acclimated to handling and head fixation.

Mice were then implanted with chronic Neuropixels 2.0 probes using one of 4 possible implants (see section ‘Implants’ below). Briefly, implantation was performed under isoflurane (1– 3% in O_2_) anaesthesia and after injection of Colvasone and Rimadyl. The UV glue was removed, and the skull was cleaned and scarred for best adhesion of the cement. The skull was levelled, and a craniotomy was performed using a drill or a biopsy punch. Once exposed, the brain was

## Code availability

The code will be made available upon publication. covered with Dura-Gel (Cambridge Neurotech). Probes were implanted in left or right primary visual cortex (2.5 mm lateral, 3.5 mm posterior from Bregma, one probe per animal).

### Implants

For the implants, we used multiple strategies: in 4 mice we used a cemented implant (*15*) and in 7 mice we used a recoverable modular implant (*15, 16, 71, 72*). The choice of implant did not affect the results.

#### Cemented implant

Mice were implanted by holding and inserting the probes using a cemented dovetail and applying dental cement to encase the probe printed circuit board and reliably attach it to the skull. The recordings were made in external reference mode, using the silver wire or the headplate as the reference signal. The data from these four mice were already published (*15*).

#### Recoverable modular implants

Mice were implanted recoverable, modular implants, such as the “Apollo” implant (*16*) (3 mice), the “Haesler” implant (*71*) (1 mouse), or the “Repix” implant (*72*) (3 mice). The methods for each implant are described in their respective papers. Briefly, after carefully positioning of the shanks at the surface of the brain, avoiding blood vessels, probes were inserted at slow speed (3–5 µm/s). Before surgery, the probes were coated with fluorescent dye DiI (ThermoFisher) by either manually brushing each probe with a droplet of DiI or dipping them in directly in DiI, for histological reconstruction. Once the desired depth was reached (optimally, just before the docking module touched the skull), the implant was sealed using UV glue, then covered with Super-Bond polymer, ensuring that only the docking module was cemented. After finishing all recording sessions, the probes were explanted and cleaned before reusing. The recordings were made in external or internal reference mode, using the headplate as the reference signal.

### Anatomical reconstruction

We imaged the brains using serial section (*73*) two-photon (*74*) tomography. Our microscope was controlled by ScanImage Basic (Vidrio Technologies) using BakingTray (*75*) (github.com/SWC-Advanced-Microscopy/BakingTray). Images were assembled using StitchIt (*76*) (github.com/SWC-Advanced-Microscopy/StitchIt). Brains were aligned to the Allen Brain Atlas (*77*) and probe location was checked using brainreg (*78*) or through custom software (www.github.com/petersaj/AP_histology). This showed that most recordings were in area VISp, and partially in high-visual areas. The exact location of the probe in visual cortex did not affect the results so we pooled all visual areas together under the name of visual cortex (VIS).

### Data acquisition and processing

Data were acquired using SpikeGLX (billkarsh.github.io/SpikeGLX/), and each session was spike-sorted with Kilosort2.5 (*15*). Well-isolated units were selected using Bombcell (*79*) (github.com/Julie-Fabre/bombcell; using parameters defined bc_qualityParamValuesForUnitMatch.m). To track neurons across days, we used UnitMatch (*17*), which uses the waveform of individual neurons within each recording to compute a pairwise probability of matching across all pairs. We used the “intermediate” level of conservativeness to determine whether units were tracked across days. Neurons were categorized as “Narrow Spiking” if their waveform duration was lower than 0.4 ms, and “Wide Spiking” otherwise.

### Stimuli

In each session, mice were presented with 112 natural images, repeated 5 times in a random order and separated by a static grey screen. The images were local contrast-normalized and were the same as in Ref. (*15*). Depending on the protocol, images were presented for 1s with fixed 2s inter-trial intervals, or for 0.5s with fixed 0.8s inter-trial intervals. “Spontaneous” periods were defined as all time-periods without a stimulus on the screen, excluding a 0.3s window after the offset of each stimulus. Visual stimuli were presented through three displays (Adafruit, LP097QX1), each with a resolution of 1024 × 768 pixels. The screens covered approximately 270 × 70 degrees of visual angle, with 0 degrees being directly in front of the mouse, and images were presented on the front and contralateral screens. The screens had a refresh rate of 60 frames per second and were fitted with Fresnel lenses (Wuxi Bohai Optics, BHPA220-2-5) to ensure approximately equal luminance across viewing angles.

### IBL dataset

We used Neuropixels recordings in the mouse brain performed in our consortium of laboratories as detailed elsewhere (*29*) (revision: 2024-05-06). Specifically, we focused on 419 sessions that contained a period of spontaneous activity, where the screen was grey. These recordings, using 644 probes, spanned 230 regions, totalling 54,553 neurons.

## Data analysis

### Stimulus-evoked activity

#### PSTHs

To compute the peri-stimulus time histograms (PSTHs), we computed for each neuron the number of spikes within 1ms time bins ranging from 25 ms before to 150 ms after the stimulus onset. We then temporally smoothed the PSTHs with a 30 ms-wide gaussian window (σ = 5.8 ms).

#### NMF model

To explain the response of single neurons across images, we performed Non-negative Matrix Factorization (NMF) on the response matrix of individual neurons *R* of elements *R*_*t,i*_ (number of time points x number of images) using a range of factors from 0 to 4 which defined the rank of our prediction:

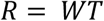

where *W* contains the tuning components (size number of factors x number of images) and *T* the temporal components (size number of timepoints x number of factors). For example, the rank-0 model consisted simply in a uniform firing rate, the rank-1 model consisted in a single time course multiplied with a single tuning profile, and more generally the rank-n model contains n time courses associated with n tuning profiles. To measure the goodness of the model, we computed the coefficient of determination R^2^ between the prediction and the test set. To compute the R^2^ across models (**Fig. 1**I and J), we used a leave-one-out cross-validation procedure where four repeats were used for training and one for testing. In **Figure 1**G and H, we used half of the repeats to plot the neurons’ response, and the other half to compute the model.

### Population coupling

We performed the same analyses on both datasets.

#### Spike-triggered population rate

To measure population coupling, we first binned the firing rate of neurons in 1ms time bins, focusing on spontaneous periods, obtaining the raw firing *f*_*i*_(*t*) of neuron *i* across time. We then computed the spike-triggered population rate (stPR, *c*_*i*,τ_), similarly to Ref. (*25*) and defined as:

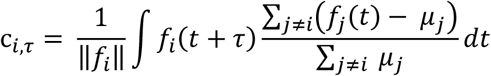

where τ represents the time lag between the neuron and the population, µ_*i*_ the average firing rate of neuron *i*, ‖*f*_*i*_‖ the number of spikes it fired, and *N* the total number of neurons in the recording. Note that compared to Ref. (*25*), we normalized by the average firing rate, and time-lagged the analysis. Importantly, the population coupling is uncorrelated with the neuron’s firing rate (*25*). Moreover, this definition is such that a negative delay corresponds to the neuron leading the population.

To focus on 10-100 ms long sequences of activity, we computed coupling within a [-80 80] ms ([-100 100] ms for the IBL dataset) window, and low-passed filtered the coupling with a 20 Hz cutoff.

#### Coupling strength

The strength of the population coupling was defined at lag 0 (*c*_*i*,0_). To summarize the overall level of coupling strength for each recording and brain area, we computed the median population coupling strength across all neurons within that brain area (**Fig. 4**E and H, **Fig. 5**G, H and L).

### Sequences

We defined a neuron’s delay by computing its centre of mass *d*, defined as follows:

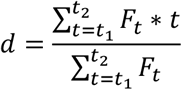

Where *F*_*t*_ is the metric (firing rate, population coupling) at time *t*, within the time window [*t*_1_, *t*_2_], normalized between 0 and 1.

#### Stimulus-evoked sequences

To compute each neuron’s delay in response to individual images, we measured the centre of mass of the PSTH averaged over the repeats of that image within a “responsive window”. The “responsive window” was defined as the timepoints within 20 to 150 ms after stimulus onset where the firing rate of the neuron was at least two standard-deviations above its baseline. When no timepoints were filling this criterion, the centre of mass was set to *nan*.

Similarly, to compute each neuron’s average delay across all images, we performed a similar analysis on the time course component given by the rank-1 model.

#### Coupling delay

To compute each neuron’s delay during spontaneous activity, we computed the centre of mass of their low-passed stPR. To visualize the sequences, we plotted a heatmap of the stPR averaged across repeats, and displayed the coupling delays for both the first and the second half of the recording, sorted by the first half.

#### Sequence width

The sequence width was defined as the difference between the 10^th^ and the 90^th^ percentiles of the coupling delays across neurons.

### Reliability, similarity and stability

#### Reliability

To measure the reliability of the various metrics (coupling strength, delay, etc.), we split all recordings into two halves and performed the analysis on both halves. For stimulus-evoked responses, we used odd and even repeats of the same stimuli as the two halves. For population coupling strength or delay, we artificially cut the recording halfway. We then defined reliability as the correlation of the metric across the two halves.

#### Similarity

To measure the similarity *s* of metrics *x* and *y* across conditions (e.g. across stimulus condition, across days), we used noise-corrected correlation (*80*). In short, noise-corrected correlation normalizes the z-transformed average correlation of the metric across conditions by the reliability of the metric within condition:

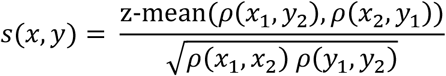

where ρ denotes the Pearson correlation coefficient, and *x*_1_, *x*_2_ and *y*_1_, *y*_2_ are the vectors of metric *x* and *y* for the two halves of the data. The function z-mean computes the average of the z-transformed correlation coefficient before inverse z-transforming it:

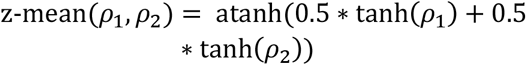

To measure the similarity between evoked and spontaneous sequences, we used recordings where at least 10 neurons of each type were present. We computed the similarity of the centres of mass of the spontaneous population couplings and the centres of mass of the time course given by the rank 1 model.

#### Stability

To measure the stability of a metric across days, we simply computed the similarity of the metric across pairs of recordings. We selected pairs of recordings where at least 15 neurons were tracked, and selected animals with at least 5 recordings, with at least one pair of recordings more than 15 days apart, yielding 4 animals. To measure the stability of evoked responses (time course and tuning), we focused on neurons that were visually responsive, as defined by being better predicted by a model with a rank > 0 rather than its average firing rate, as defined by its cross-validated R^2^.

The stability was then estimated performing a robust regression using the bisquare weighting function (*fitlm* in Matlab, with option ‘RobustOpts’ on). The average of the intercept and slope across mice was then used to show the average fit.

### Reliable variance partitioning

To measure the contribution of layers to the distributions of population coupling strengths or delay we observed in cortex, we partitioned the total reliable variance (*v*_*tot*_) of these distributions into two components, the variance across layers (*v*_*across*_), and the variance within layers (*v*_*within*_):

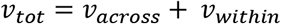

For a metric of interest with two repeats *x*_*n*,1_ and *x*_*n*,2_ for neurons *n*, we defined the reliable variance as the unnormalized covariance of the two repeats across neurons. Thus, the variance within layers is given by:

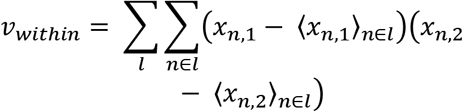

Where 〈*x*_*n*,1_〉_*n*∈*l*_ denotes the mean across neurons belonging to layer *l*, i.e., layer 1, 2/3, 4, 5, 6a and 6b. Conversely, the variance across layers is given by:

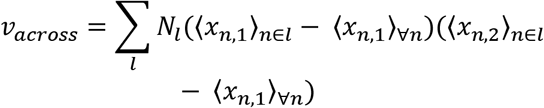

Where *N*_*l*_ is the number of neurons in layer *l*. The proportion of reliable variance not explained by layers, *p*, is then given by:

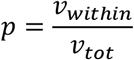

Layers with 1 neuron were excluded since they cannot help disentangling the contribution of neuron identity vs. layer.

### Cross-area analysis

#### Cross-area coupling

To compute cross-area coupling, we simply computed the time-lagged coupling of each neuron from one area (the ‘target’ area) with the average population rate of the other area (the ‘source’ area). We then computed the coupling strength and delay of each neuron of the target area relative to the source area.

We then obtained, for each metric of interest, a matrix of size (number of areas x number of areas) that was sparsely populated, depending on the available paired recordings. We then defined for each metric two summary statistics for each brain area: the average metric as a source (averaging over targets, excluding the area itself) and the average metric as a target (averaging over sources, excluding the area itself).

#### Connectivity and distance

We related the cross-area sequences to two anatomical metrics: axonal connectivity (axonal projection tracing (*29*)) and cartesian distance (inverse Euclidean distance between centroids of region pairs). To compute the connectivity between brain areas, we directly loaded the 41586_2014_BFnature13186_MOESM70_ESM.csv from Ref. (*32*) and extracted the quantitative projection signals for each pair of areas. To compute the Euclidean distance between brain areas, we used the inverse distance between the centroids of each area, as in Ref. (*29*). To determine the significance of the correlation between the connectivity and the coupling delays reliability between areas, we used a Mantel test. We computed the Spearman coefficient for the real data and for 10,000 permuted matrices (keeping the same permutation order for rows and columns). The correlation coefficient for the real data was above that of all permutations.

### Inclusion criteria for the IBL dataset

The IBL dataset contains recordings on both hemispheres. For within-region calculations (e.g., coupling delay reliability with their own population rate), we treated each hemisphere as an independent recording. For cross-regions calculations, we focused on a single hemisphere, the right one for the cerebellum and the left one for the rest of the brain, which contained most of the recordings.

To obtain the average values for each metric and each brain area across the IBL dataset, we selected recordings with more than 5 neurons in each area of interest. For example, for cross-area analysis, we focused on paired areas where more than 5 neurons were recorded in both areas.

For within-area coupling (**Fig. 4**), we further selected areas with at least 5 recordings. For cross-area coupling, we selected pairs of areas with at least 2 recordings for **Fig. 5**G to J but took all pairs for **Fig. 5**L to M. This was because our conclusions do not focus on specific pairs of regions, but on the global reliability of coupling strengths and delays across the brain.

---

**Supplementary Figure 1.**
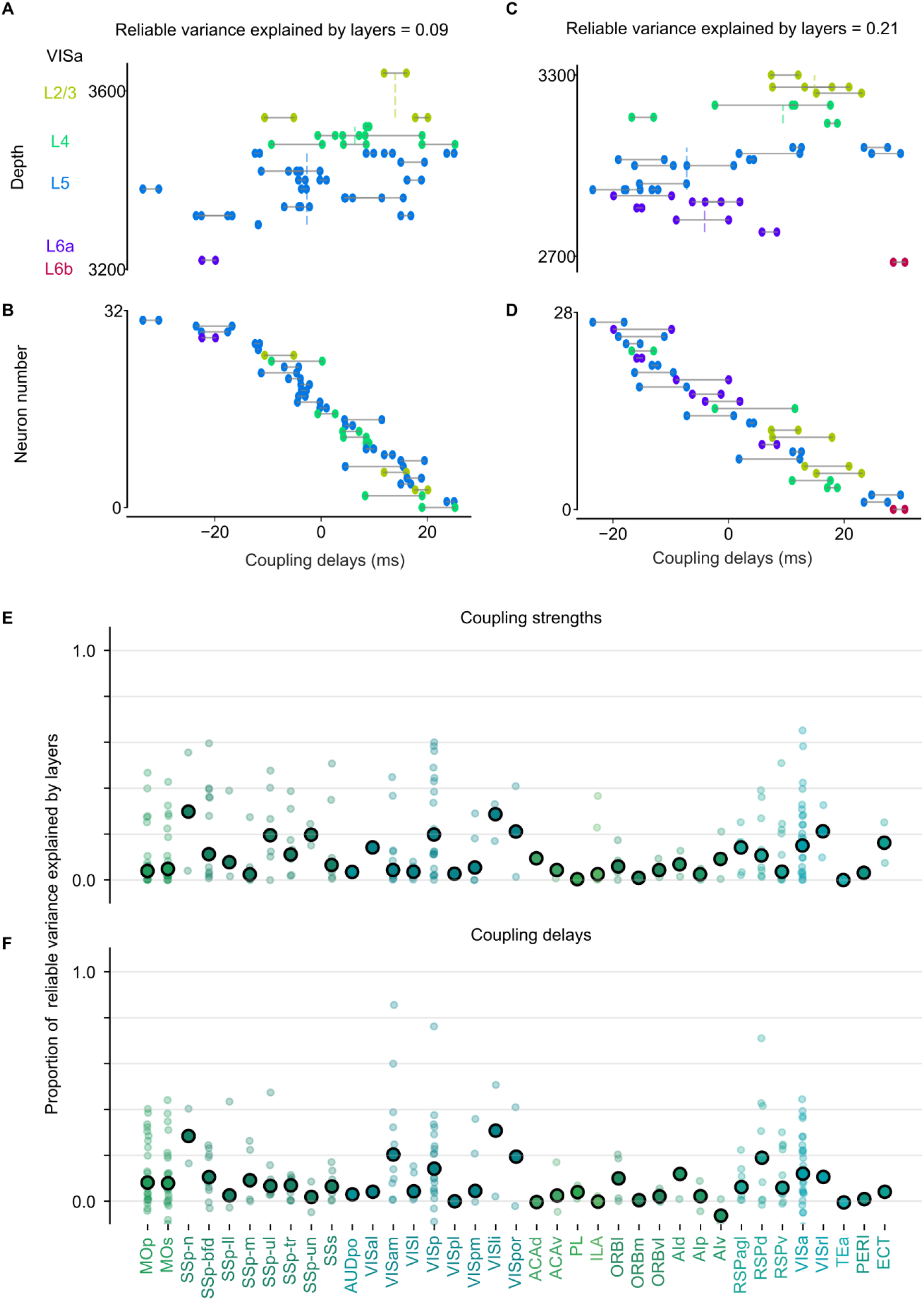
Coupling strengths and delays are largely independent of the layer structure in cortex. (**A**) Relationship between the coupling delays of individual neurons (computed on 1^st^ and 2^nd^ half of the recording, as shown by the connected two dots) as a function of their depth on the probe in an example recording in VISa. The colours indicate the layer to which the neurons belong. (**B**) Ordering of the same neurons as a function of their coupling delay in the first half. Only 9% of the reliable variance was explained by the layer structure. (**C**) and (**D**) sane as (A) and (B) for another recording, where layers explain 21% of the reliable variance. Even in that case, large variations of coupling delay are observed within layers. (**E**) Summary across all recorded cortical regions of the reliable variance of the distribution of coupling strengths not explained by layers, showing individual recordings (*small dot*) and the median across recordings (*large dot*) for each region. (**F**) Same as (E), for coupling delays. Values can be above 1 because of the cross-validation.

